# The genetic basis of novel trait gain in walking fish

**DOI:** 10.1101/2023.10.14.562356

**Authors:** Amy L Herbert, Corey AH Allard, Matthew J McCoy, Julia I Wucherpfennig, Stephanie P Krueger, Heidi I Chen, Allex N Gourlay, Kohle D Jackson, Lisa A Abbo, Scott H Bennett, Joshua D Sears, Andrew L Rhyne, Nicholas W Bellono, David M Kingsley

**Affiliations:** Department of Developmental Biology, Stanford University School of Medicine, Stanford CA 94305 USA; Department of Molecular and Cellular Biology, Harvard University, Cambridge MA 02138 USA; Department of Pathology, Stanford University School of Medicine, Stanford CA 94305 USA; Roger Williams University, Bristol, RI 02809 USA; Marine Biological Laboratory, Woods Hole, MA, 02543 USA; Howard Hughes Medical Institute, Stanford University School of Medicine, Stanford CA 94305 USA

## Abstract

A major goal in biology is to understand how organisms evolve novel traits. Multiple studies have identified genes contributing to regressive evolution, the loss of structures that existed in a recent ancestor. However, fewer examples exist for genes underlying constructive evolution, the gain of novel structures and capabilities in lineages that previously lacked them. Sea robins are fish that have evolved enlarged pectoral fins, six mobile locomotory fin rays (legs) and six novel macroscopic lobes in the central nervous system (CNS) that innervate the corresponding legs. Here, we establish successful husbandry and use a combination of transcriptomics, CRISPR-Cas9 editing, and behavioral assays to identify key transcription factors that are required for leg formation and function in sea robins. We also generate hybrids between two sea robin species with distinct leg morphologies and use allele-specific expression analysis and gene editing to explore the genetic basis of species-specific trait diversity, including a novel sensory gain of function. Collectively, our study establishes sea robins as a new model for studying the genetic basis of novel organ formation, and demonstrates a crucial role for the conserved limb gene *tbx3a* in the evolution of chemosensory legs in walking fish.

## Main

Understanding the molecular underpinnings of evolutionary change is a key goal in biology. Recent studies have identified multiple examples where discrete loci control much of the variation in particular traits^1^ including loss of bristles in flies^2^, loss of pigment in cavefish^3^, loss of legs in snakes^4^, and reduction of armor plates and pelvic spines in sticklebacks^5,6^. Frequently, key evolutionary loci encode essential developmental regulators, with *cis-*regulatory mutations making it possible to avoid negative pleiotropy and confine major effects to particular tissues^2,4,6–10^. However, many of the best understood examples of evolutionary change involve trait loss. Whether the evolution of gained traits involve similar types of key developmental genes and *cis-*regulatory changes is not yet clear, particularly in vertebrates. Moreover, while computational studies have begun to examine the molecular basis of novel vertebrate traits, such as evolution of flight in birds and mammals^11^, headgear in ungulates^12^, or echolocation in bats and cetaceans^13^ it has usually not been possible to functionally test the hypotheses generated from comparative genomic studies in the same species. To address these issues, we established sea robins as a new model system for studying the development and evolution of novel traits in vertebrates. Sea robins are marine fish that exhibit striking evolutionary novelties, including enlarged, winglike pectoral fins, and six prominent detached “legs,” which provide sensory and locomotor capabilities to walk, dig, and detect food on the ocean floor^14,15^ (**Fig. 1a**). Sea robin leg development is a particularly salient example of trait gain because 1) it involves major alterations to skeletal patterning within a well-known limb bud/appendage development structure and 2) it is accompanied by extensive increases in the musculature and sensory and motor neuronal systems, including formation of novel enlarged lobes in the central nervous system^1^. Here, we develop two readily available sea robin species (*Prionotus carolinus* and *Prionotus evolans*) that exhibit notable differences in pectoral fin size, pigmentation pattern, and leg structure (**Extended Data Fig. 1**) for use in a laboratory setting, and then use a combination of genomic and functional studies to probe the origins of evolutionary novelty in vertebrates.

**Fig. 1:**
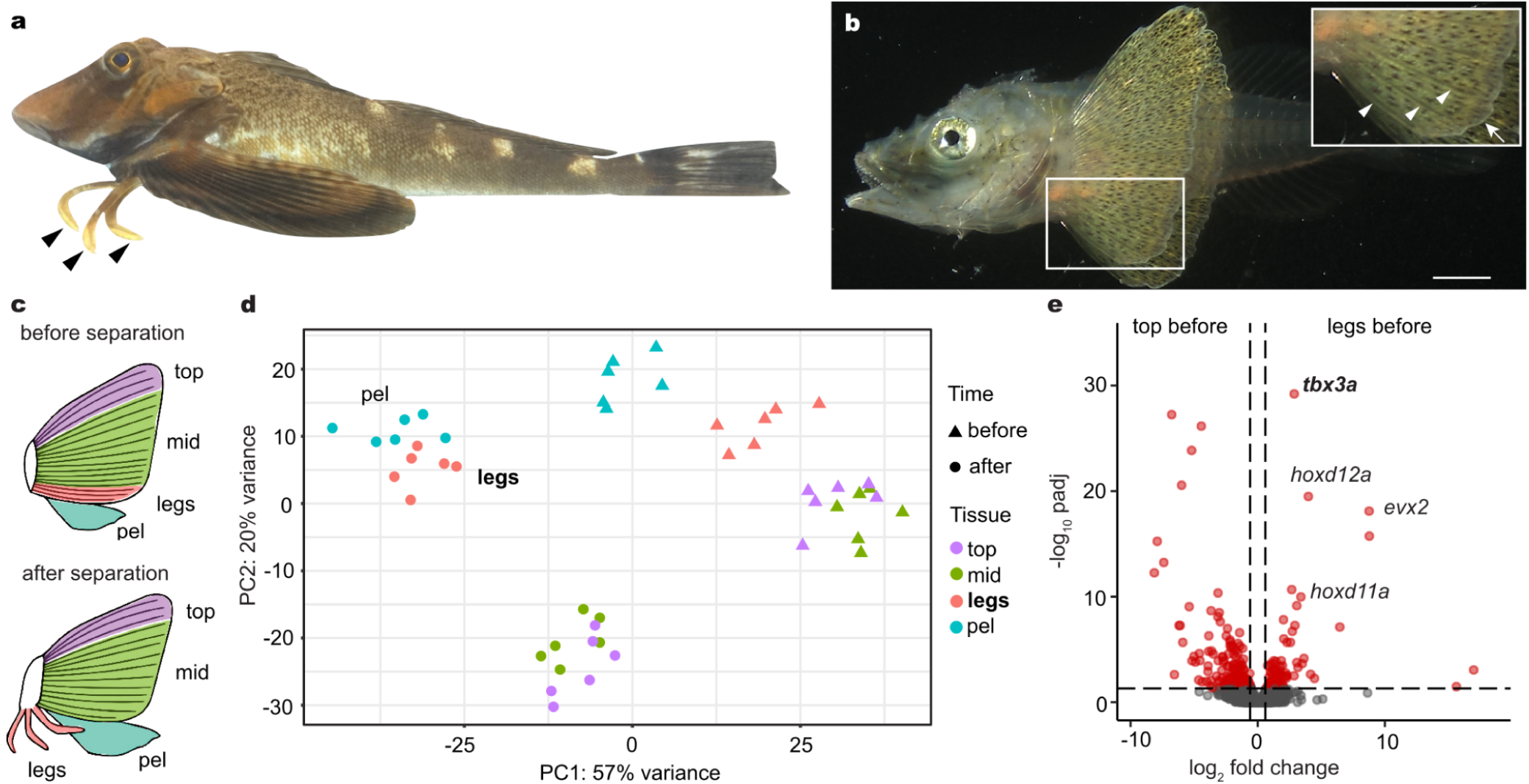
Sea robin leg development and molecular profiling. **a,** Sea robins are fish that exhibit novel leg-like structures (arrowheads). **b,** Legs develop from the bottom three fin rays initially connected to the rest of the pectoral fin. White arrowheads point to three connected fin rays and arrow points to fin webbing. **c**, Diagrams of fin tissues collected for RNA-seq at two time points, before and after leg separation. The pectoral fin was dissected into the top three fin rays (top), the middle part of the pectoral fin (mid) and the bottom three rays or legs (legs), depending on the time point. The pelvic fin (pel) was collected in its entirety. **d**, A PCA plot of all tissue samples shows legs beginning to cluster with the pelvic fin before separation. After separation, legs cluster with pelvic fins instead of the other pectoral fin components. **e**, Volcano plot showing upregulated expression of *tbx3a*, *hoxd12a*, *evx2*, and *hoxd11a* in future legs compared to the top portion of the pectoral fin before leg separation. Exact *P-adjusted* values: *padj* = 5.54e-31 (**e**, *tbx3a*), *padj* = 7.19e-20 (**e**, *hoxd12a*), *padj* = 1.26e-18 (**e**, *evx2*), *padj* = 1.68e-10 (**e**, *hoxd11a*). N = 6 animals before leg separation and N = 6 animals after leg separation. Scale bar, 1 mm (**b**).

### Developmental mechanisms of leg development

To utilize sea robins as an experimental model, we established *in vitro* fertilization and a sea robin culturing system to generate embryos and grow larvae for developmental studies (see Methods). Consistent with previous morphological studies^17,18^, sea robin “legs’’ develop from the three ventral most pectoral fin rays, which initially form within the pectoral fin (**Fig. 1b**), and then separate from the rest of the fin rays during development. To molecularly characterize leg development, we sequenced, assembled, and annotated the genome of the Northern sea robin *P. carolinus*. We then performed tissue specific RNA-sequencing (RNA-seq) before and after the leg rays separate, and compared expression patterns in the ventral leg rays (legs), to the developing pectoral and pelvic fins of *P. carolinus* (**Fig. 1c**). Although originating from the pectoral fin, PCA analysis revealed that leg rays are molecularly more similar to the pelvic fin than to the other pectoral fin rays (**Fig. 1d**). Some of the top upregulated genes distinguishing developing leg rays from the rest of the pectoral fin include well known developmental transcription factors that have a role in both pectoral fin development as well as tetrapod limb development, such as *hoxd12a*, *hoxd11a* and *evx2*^19–21^ (**Fig. 1e**). We note that *hoxd12a* appeared to be the highest numbered *hoxd* paralog present in the assembled and annotated sea robin genomes, suggesting that some of the previously described limb functions of *hoxd13a* in other fish and tetrapods^9,21,22^ have likely been modified in sea robins. We also identified markers of pectoral fin rays, including *tbx15*, and the actinodin genes *and1* and *and2* (**Extended Data Fig. 2**). *Tbx15* has a role in shoulder girdle formation and is expressed in zebrafish pectoral rays^23,24^, while a*nd1*/*and2* constitute major structural components of fish fin rays. Loss of these two genes in land animals has previously been associated with the evolutionary transformation of fins to limbs in tetrapods^25^. Reduced expression of actinodin genes in sea robin legs highlights the structural differences between legs and pectoral fin rays in a walking fish lineage. The sequencing studies provide a set of marker genes that distinguish leg and fin fate, as well as candidate genes that sea robins may use to specify their novel locomotory appendages.

The top differentially expressed gene between the leg rays and other fin rays in the pectoral fin prior to leg separation was *tbx3a* (**Fig. 1e**). This gene was also the top upregulated gene in legs compared to both the top pectoral fin (*padj* = 8.97e-31) and middle pectoral fin *(padj* = 1.27e-21) after separation, suggesting that *tbx3a* may be playing multiple developmental roles. *Tbx3* encodes a T-box transcription factor and is present in a genomic cluster with *tbx5*, a master regulator of forelimb development^26^. Although *tbx3a* was duplicated in the teleost lineage that includes sea robins, we only observed differential expression of *tbx3a*, not *tbx3b,* in the fin tissues sequenced. Interestingly, humans heterozygous for mutations in *TBX3* have skeletal alterations that primarily affect structures that develop from the posterior region of the forelimb^27,28^. The preferential expression of *tbx3a* in the leg ray region of the developing sea robin pectoral fin, and the regional specificity of the skeletal alterations in humans suggested that *tbx3a* could play a critical role in leg formation.

### Functional testing of leg developmental genes

To test the functional relevance of the genes identified through RNA-seq studies, we established CRISPR-Cas9 genome editing in sea robins. We first targeted the pigmentation gene *slc24a5*, which provided a visual readout of successful editing^29^ (**Extended Data Fig. 3a**). All injected larvae showed obvious pigmentation abnormalities, and genomic amplicon sequencing showed an average of 81% mutant reads at the *slc24a5* locus, indicating that editing is highly efficient. Because genome-editing was done in the F0 generation, and the resulting fish appeared to be mosaic for multiple mutations, we will hereafter refer to CRISPR-Cas9 injected animals as crispants. We next used CRISPR-Cas9 genome editing to target genes of interest identified in our developmental RNA-seq studies. Disruption of the *hoxd12a/hoxa13a/hoxa13b* genes reduced the number of legs in sea robins (**Extended Data Fig. 3c, d, e**). Intriguingly, while disruption of *tbx3a* led to a variety of phenotypes, including angled legs and both decreases and increases in the number of legs from one to five, a notable phenotype was the development of smaller reduced legs, which frequently appeared more similar to pectoral fin rays (**Fig. 2a, b**). Notably, humans heterozygous for *TBX3* mutations exhibit similarly variable phenotypes, including both increases and decreases in number of digits, as well as alterations to direction of developing arms^27,28^. Together, our results show that multiple genomic loci can be effectively targeted in sea robins, and confirm roles for *hox* genes in determining leg number, and *tbx3a* in leg development.

**Fig. 2:**
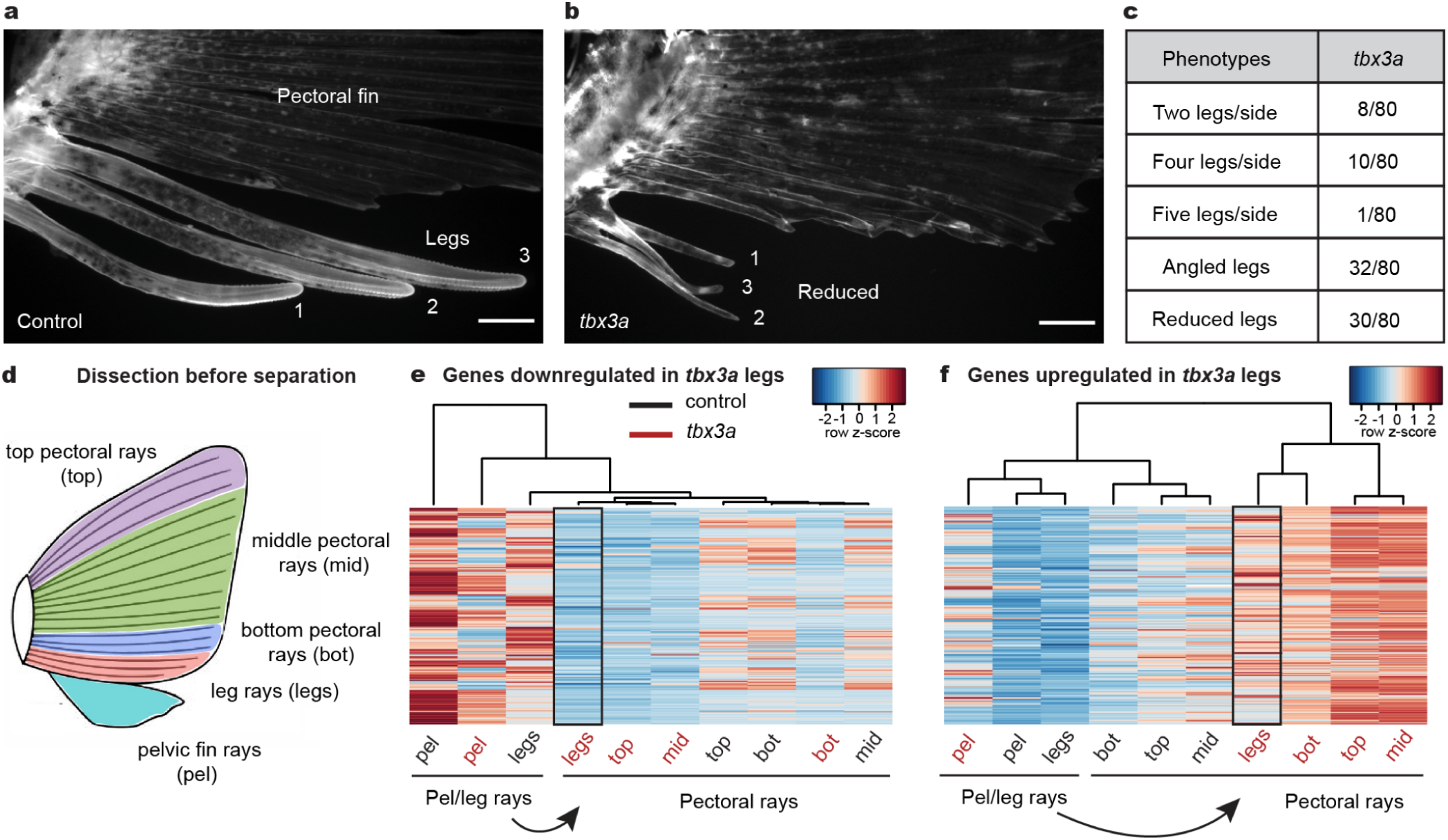
*Tbx3a* disruption alters leg number and identity. **a,** A lateral view of numbered WT legs in a control sea robin. **b**, Legs 1, 2, and 3 are reduced in a *tbx3a* crispant, and leg 1 is angled away from its usual position **c**, Quantification of crispant phenotypes in *tbx3a* injected animals (number of animals with phenotype/total number of animals phenotyped). **d**, Schematic of pectoral fin dissection into the top three pectoral rays (top), the middle pectoral rays (mid), the bottom three pectoral rays (bot) and the bottom three putative leg rays before separation (legs). The entire pelvic fin (pel) was collected in its entirety. **e**, A heatmap of genes downregulated in *tbx3a* crispant legs compared to control legs (*padj* < 0.1). **f**, A heatmap of genes upregulated in *tbx3a* crispant legs compared to control legs (*padj* < 0.1). N = 8 control animals and N = 10 *tbx3a* crispants. Scale bars, 1 mm (**a**, **b**).

### Functional and molecular changes in *tbx3a* targeted fish

The observation that reduced *tbx3a* legs showed morphological similarities to fin rays led us to hypothesize that *tbx3a* might be a key driver of leg specification. To understand the molecular consequences of *tbx3a* disruption in legs, we performed RNA-seq profiling of *tbx3a* crispants and control siblings before and after leg separation. When we compared the genes downregulated or upregulated in putative *tbx3a* legs before separation compared to control legs, we found that *tbx3a* legs clustered with pectoral fins instead of control legs (**Fig. 2d, e, f**). Furthermore, we found that 40% of the pectoral fin marker genes were upregulated in *tbx3a* crispant leg rays compared to control leg rays (**Extended Data Fig. 4a**) while none of these genes were upregulated in control legs compared to *tbx3a* crispants (**Extended Data Fig. 4b**). Pectoral fin markers upregulated in *tbx3a* crispants included several genes involved in pigmentation pathways, such as *pmela*, *tryp*, *slc24a5*, and *tryp1b* (**Extended Data Fig. 4c**), consistent with a previously studied role for *Tbx3* in pigmentation suppression in horses^30^. Overall, these results demonstrate that disruption of *tbx3a* results in upregulation of pectoral fin markers prior to leg separation, indicating that leg rays become more similar to fins in the absence of *tbx3a*.

We performed a similar molecular analysis on individual legs from *tbx3a* crispants and uninjected controls after leg separation (**Extended Data Fig. 5a**). Again, approximately 20% of pectoral fin marker genes were also upregulated in *tbx3a* crispant leg 3 (which had the strongest pectoral fin phenotype) compared to control leg 3 (**Extended Data Fig. 5b**). In contrast, only 0.9% of pectoral fin marker genes were upregulated in uninjected leg 3 compared to *tbx3a* crispant leg 3 (**Extended Data Fig. 5c**). We note that some *tbx3a* crispants had two legs instead of three, and one crispant had four legs, an example of variability in the fish system which resembles both the increases and decreases in digit number previously reported in human Ulnar-mammary patients^28^. Comparison of genes upregulated in pectoral fins and leg 3 highlights the 45 genes driving *tbx3a* crispant leg 3 to become more similar to the pectoral fin (**Extended Data Fig. 5d**). We also identified upregulation of specific pectoral fin genes in leg 3 of *tbx3a* crispants, including *and1* and *tbx15* (**Extended Data Fig. 5e**). Although not significant, we also saw higher expression of *and2* counts in *tbx3a* crispant leg 3 (**Extended Data Fig. 5e**). Upregulation of these fin ray genes in *tbx3a* crispant legs thus supports a key role for *tbx3a* in promoting a leg versus pectoral fin ray fate.

To further investigate how loss of *tbx3a* might functionally impact sea robins, we examined the striking dorsal accessory spinal lobes of the sea robin nervous system. These macroscopic CNS lobes are elaborations of the dorsal horn of the spinal cord that are organized in a one-to-one relationship with legs and are thought to play a role in leg function^16,31^ (**Fig. 3a**). We found that the CNS lobes of sea robin *tbx3a* crispant animals were significantly reduced compared to the lobes of control siblings (**Fig. 3b, c**). In a companion study, we showed that *P. carolinus* uses legs as sense organs to efficiently find and unearth buried food items (**Fig. 3d**). We used this behavioral assay to test the functional consequences of *tbx3a* disruption and found that *P. carolinus tbx3a* crispants exhibited reduced ability to localize and uncover buried mussels, close to levels of the non-digging species *P. evolans* (**Fig. 3e**). Thus, disruption of *tbx3a* had a profound effect on sea robin behavior, consistent with the altered legs and reduced neural elaboration.

**Fig. 3:**
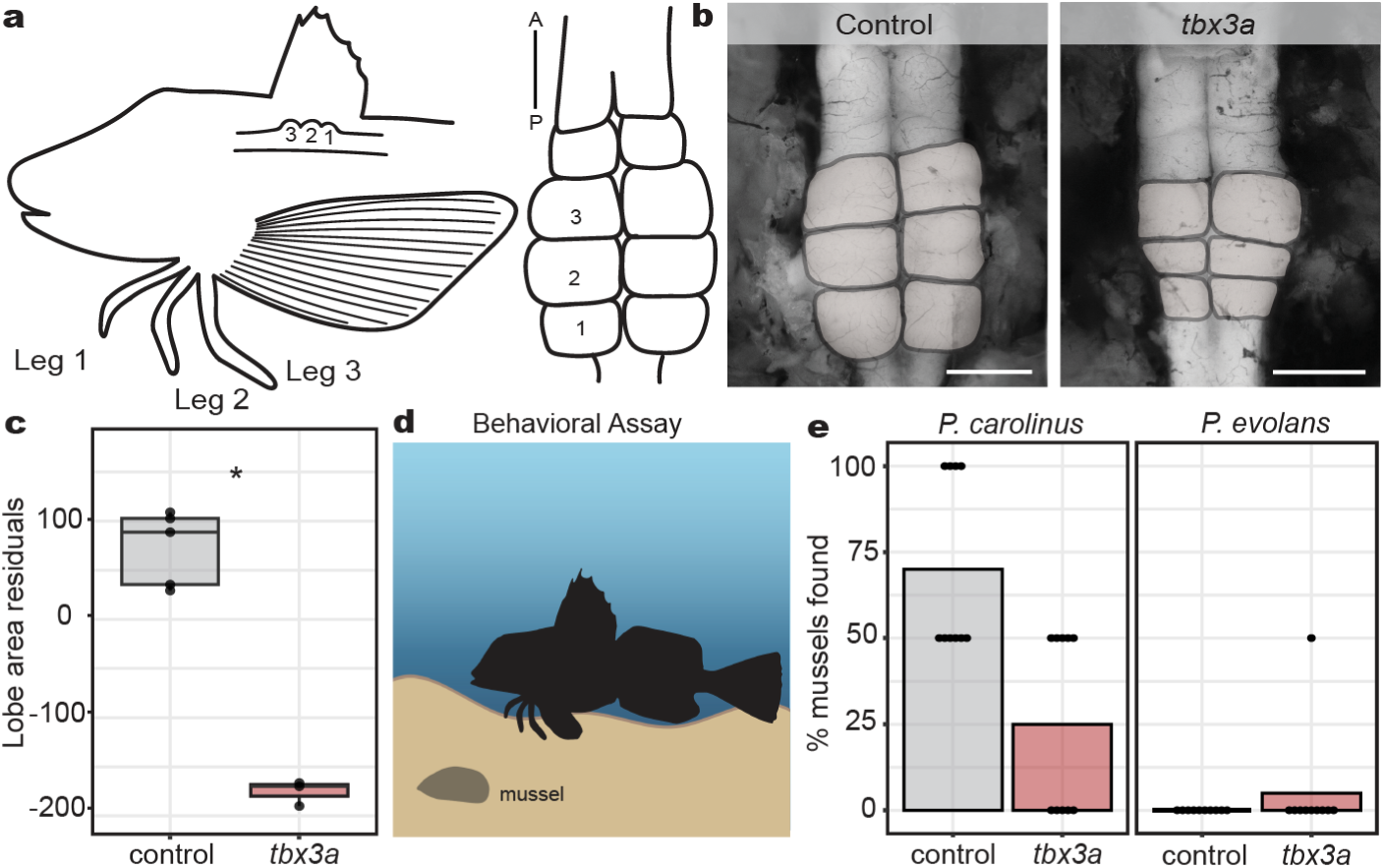
Effects of *tbx3a* mutation on leg neural architecture and behavior. **a**, Diagram of the 1:1 innervation relationship between sea robin legs and CNS lobes. **b**, Representative examples of control and *tbx3a* crispant lobes (lobes pseudo colored in gray). **c**, Lobe area was reduced in *tbx3a* crispants as measured by regressing against standard length (N = 5 control animals and N = 3 *tbx3a* crispants, *P* = 0.04 by Wilcoxon rank sum test). **d**, Schematic of behavioral assay used to test sensory behavior of sea robins (created with biorender). **e**, Digging behavior was reduced in *tbx3a* crispants versus control *P. carolinus*, closer to levels of non-digging *P. evolan*s controls that were unaffected by *tbx3a* disruption. Two mussels were buried per trial. Discovery of both mussels resulted in a score of 100% while uncovering one mussel was scored as 50%. N = 10 trials across 6 animals per genotype used in analysis. Scale bars, 1 mm (**b**, both images).

### *Cis-* and *trans-*regulatory control of species-specific differences

The two sea robin species *P. carolinus* and *P. evolans* exhibit striking phenotypic differences, such as pectoral fin size, coloration, and epithelial papillae and increased digging abilities in *P. carolinus* legs, making them an excellent model for studying the molecular underpinnings of species-specific differences. To develop this comparative model, we sequenced, assembled, and annotated the genome of *P. evolans* to analyze the molecular basis of leg formation and function. Furthermore, using *in vitro* fertilization, we successfully crossed *P. evolans* females and *P. carolinus* males to produce viable hybrid animals that survived through the leg separation stage (**Fig. 4a**). To analyze the genetic basis of diversification of leg function, we collected legs and fins from individual species and hybrid animals before and after leg separation. We then used RNA-seq to identify differentially regulated genes between the two sea robin species, and genes with allele-specific expression differences in F1 hybrids. This experiment makes it possible to identify the relative contribution of *cis-* and *trans-*acting changes to species-specific gene expression differences, as *cis-*regulatory changes should persist in F1 hybrids, while differences due to *trans-*regulatory effects should disappear when both genomes are present in the same *trans-*acting environment^32^ (**Fig. 4b**). Comparisons between the two species showed a high percentage of differentially expressed genes, consistent with differences in morphology and function (**Fig. 4c**), and a large fraction of these genes also showed allele-specific expression in the interspecific hybrids, (adjusted R^2^ = 0.4553, *p* < 2.2e-16, **Fig. 4c** and **Extended Data Figs. 6 and 7**). Overall, these data suggest that the majority of the leg and fin expression differences observed between the two species are regulated in *cis-*. However, a substantial number were regulated in *trans-,* or by combinations of both *cis-* and *trans-*regulation, indicating a potential role for *trans-*regulatory changes in driving the evolution of sea robin traits.

**Fig. 4:**
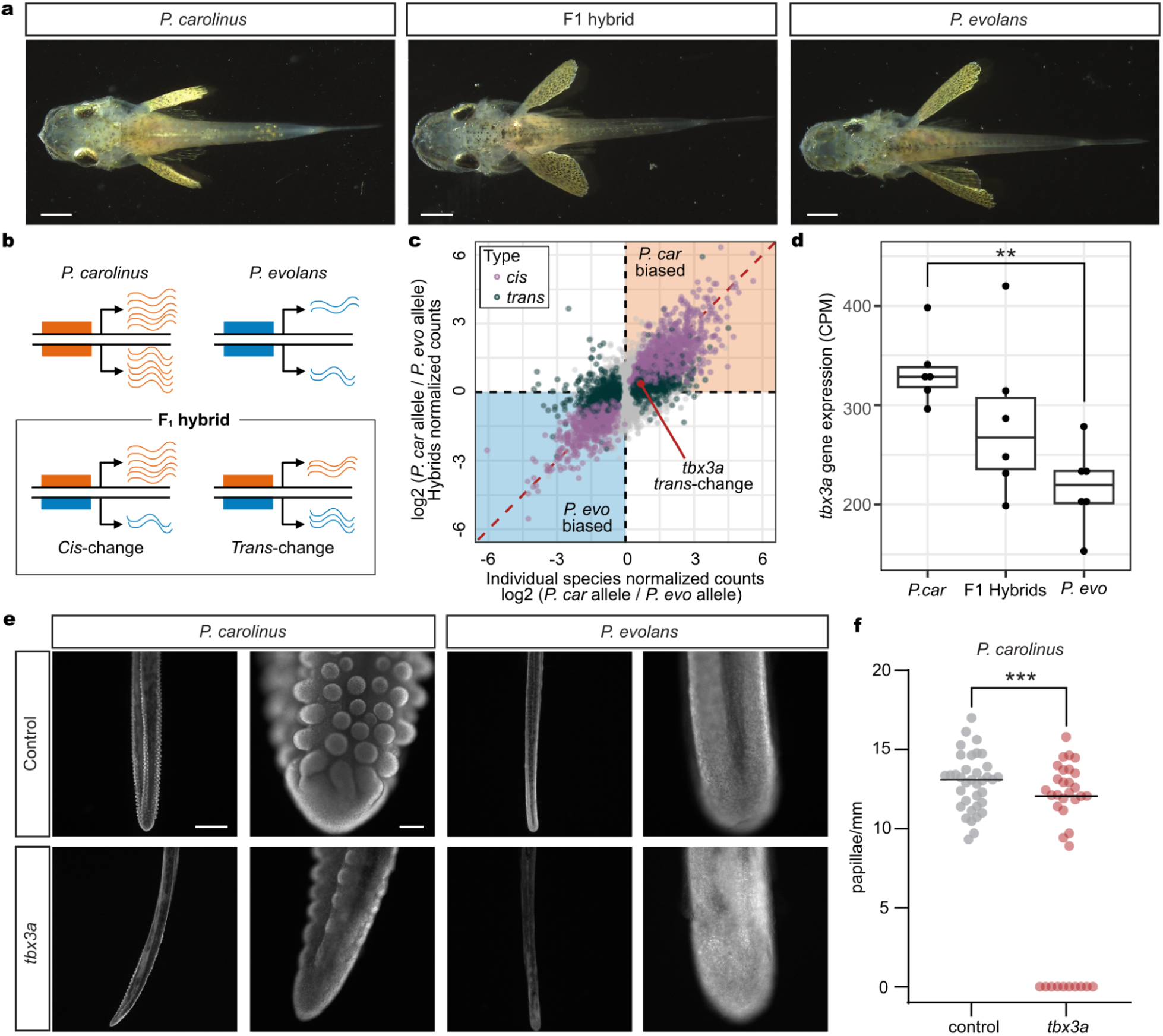
Hybrids reveal species-specific gene regulation. **a,** Dorsal views of *P. carolinus*, F1 hybrid, and *P. evolans* animals. **b**, A diagram shows how *cis*- and *trans*-regulation can be delineated by comparing gene expression differences in two separate species to allele-specific expression differences in F1 hybrid animals. **c**, Scatterplot showing gene expression differences between the two species (x-axis; N = 6 *P. carolinus*, N = 5 *P. evolans*) and the differences between the parental alleles in the F1 hybrid animals (y-axis; N = 6 F1 hybrids) in legs after separation. Genes are labeled as *cis*-regulated (purple; differential gene expression *padj* < 0.05 and allele-specific expression *padj* < 0.05) or *trans*-regulated (dark green; differential gene expression *padj* < 0.05 and allele-specific expression *padj* ≥ 0.05). Identity line is shown in red. **d**, RNA expression of *tbx3a* in counts per million (CPM) in *P. evolans,* F1 hybrids, and *P. carolinus* in legs after separation. Expression is higher in *P. car* compared to *P. evo* (log2 fold change = 0.67, *padj* = 0.0027). **e**, Control *P. carolinus* legs show abundant papillae (N = 5/5) that were reduced by *tbx3a* disruption (N = 2/3) while *P. evolans* legs lacked papillae in both control (N = 3/3) and *tbx3a* disrupted conditions (N = 3/3). **f**, Quantification of papillae density in *P. carolinus* controls (N = 6) and *tbx3a* crispants (N = 6). Exact P value: *p* = 0.0005, unpaired t-test with Welch’s correction. Scale bars, 1 mm (**a**); 1 mm, 100 μm (**e**).

Gene expression differences were observed in both directions between species. For example, the *aldh1a2* gene, which is involved in retinoic acid synthesis, was expressed at higher levels in the pectoral fins of *P. evolans* than *P. carolinus* (log2 fold change = 2.58*, padj* = 0.00087), and this difference was largely maintained in hybrids (log2 fold change = 2.08, *padj* = 0.0079)*. Aldh1a2* (or *raldh2*) has a role in outgrowth of limb appendages, and has been hypothesized to contribute to both expansion of pectoral fins in flying fishes^33^, and reduction of wings in the flightless emu^34^. Because *P. evolans* has substantially larger pectoral fins than *P. carolinus,* increased expression of the *P. evolans aldh1a2* allele, both in separate species and in F1 hybrids, suggests that *cis-*regulatory changes to *aldh1a2* may contribute to repeated evolution of forelimb size proportions across multiple fish and bird species.

In contrast, *tbx3a* was expressed at higher levels in the legs of *P. carolinus* than *P. evolans* (**Fig. 4d**), with analysis in F1 hybrids indicating that the expression difference arises primarily from *trans-*acting changes. To test for possible species-specific roles of *tbx3a*, we also targeted the gene in *P. evolans*. We found developmental leg phenotypes similar to those observed in *P. carolinus* (**Extended Data Fig. 8**), suggesting that *tbx3a* plays an early role in leg formation in both species. However, *P. carolinus* sea robins also characteristically develop wider, more robust legs than *P. evolans* (**Extended Data Fig. 1)**, as well as the novel presence of numerous epithelial papillae along the surface of the legs, which mediate increased digging and sensory capabilities (*Allard et al., 2023, companion manuscript*). Interestingly, *Tbx3* was recently hypothesized to play a role in the formation of epithelial papillae in the stomachs of ruminants^35^. Moreover, while there was a significant difference in *tbx3a* expression between the two species, F1 hybrids had an intermediate level of *tbx3a* transcripts compared with *P. carolinus* or *P. evolans* (**Fig. 4d**). This fits with our observation that hybrid animals have an intermediate papillae phenotype (**Extended Data Fig. 9a, b, c**). Notably, *P. carolinus tbx3a* crispant legs showed a significant reduction in papillae formation (**Fig. 4e, f, Extended Data Fig. 9**), with thinner legs and frequently lacked papillae altogether (**Fig. 4f)**. As expected, both *P. evolans* control and *tbx3a* crispant animals did not exhibit papillae. We cannot yet distinguish whether papillae changes arise from differences in early leg specification or from a direct role of *tbx3a* in later papillae development, but these results show that *tbx3a* function is required for a species-specific gained trait in sea robins.

Walking on legs has evolved multiple times in different organisms, including in sea robins. Here, we show that leg formation is driven by key developmental transcription factors such as *hox* paralogs and *tbx3a*. In particular, *tbx3a* appears to play a prominent role in driving specification of legs, which emerge on the post axial side of the pectoral fin. Similar expression profiles of *TBX3* in sea robin legs and human forelimbs likely reflects a conserved role in patterning structures within the postaxial zone of the forelimb bud. Leg formation in a walking fish thus utilizes an ancient pathway for regionalization of skeletal structures, illustrating how both disease phenotypes and evolutionary innovations can emerge as alternative manifestations of a key developmental control gene. Our hybrid studies indicate that many species-specific differences in leg and pectoral fin expression of *P. carolinus* and *P. evolans* have arisen through *cis-*regulatory differences, consistent with studies positing that *cis-* changes are the predominant mode of regulatory change in evolution^36^. Interestingly, our data also suggest a role for *trans*-regulatory control of *tbx3a* in novel papillae formation in sea robins, indicating that in some cases, *trans-*regulatory changes across species can drive evolutionary trait gain. The methods established here provide a new model for combining genomic and functional studies of evolutionary innovations within the larger group of sea robin relatives, which encompass a huge diversity of phenotypes, including changes to pectoral and pelvic fins, evolution of new lobes in the nervous system, development of armor and venom, and evolution of dramatic coloration patterns. Forward and reverse genetic approaches in this clade will be a powerful combination to uncover the mechanisms behind the repeated evolution of novel skeletal, sensory, and neural structures in vertebrates.

## Acknowledgements

We thank A Fire for support and helpful discussions, and V Behrens, G Roberts Kingman, and all members of the Kingsley and Bellono labs for useful input and assistance. We thank all personnel at the Marine Biological Laboratory, Roger Williams University, and Harvard University who were involved in sea robin care. In particular, we thank D Remsen, D Calzarette, E Pyszka, and N Gibbons at the Marine Biological Laboratory, A Leonardi, Z Sotero, A Grove, and L Fitzgerald at Roger Williams University, and B Walsh and P Killian from Harvard University. We thank C Albertin for providing reagents, N Ahuja for helpful conversations to develop sea robin microinjections, and J Rosenthal for helpful discussions and use of the Genome Editing Core at the Marine Biological Laboratory.

This work was supported in part by a Helen Hay Whitney Fellowship (ALH), Grass Fellowships in Neuroscience from the Marine Biological Laboratory (ALH, MJM), Early Career Whitman Fellowships from the Marine Biological Laboratory (three to ALH; two to MJM); the NIH (R35GM142697 to NWB) and the National Science Foundation (NSF-PRFB 2010728 to CAA). DMK is an investigator of the Howard Hughes Medical Institute.

## Author contributions

ALH and JIW developed hybridization and CRISPR-Cas9 editing methods for sea robins. ALH, ALR, ANG, JDS, SHB, and LAA developed aquaculture methods. ALH, MJM, JIW, HIC generated sea robin genomes. ALH and MJM performed RNA-seq and *cis-trans* analysis. ALH generated and studied crispant fish, with genotyping contributions from KDJ. CAHA, SPK, and NWB analyzed species-specific papillae, lobe, and behavioral phenotypes. ALH and DMK wrote the paper with contributions from all authors.

## Methods

### Ethical compliance and animal care

All relevant ethical regulations were followed during this study. All sea robin studies were performed according to recommendations in the Guide for the Care and Use of Laboratory Animals of the National Institutes of Health^37^. Protocols for sea robin care and use were approved by the Institutional Animal Care and Use Committees (IACUC) at the Marine Biological Laboratory (protocol numbers 17-36, 18-08C, 19-28, 20-22, 21-22, and 22-21), Roger Williams University (protocol numbers R19-07-06 and R-22-11-30), Stanford University (protocol number 32297), and Harvard University (protocol number 18-05-324-1).

### Adult sea robin care and in vitro fertilization

Adult *Prionotus evolans* and *Prionotus carolinus* were provided by the Marine Biological Laboratory (MBL) during the reproductive months of May and June. At the MBL, animals were kept in large tanks with constant ambient sea water flow and aeration by a large airstone. Animals were fed every other day. Embryos were obtained through strip spawning. In brief, eggs were obtained through manual expression of reproductive females into a large petri dish. Milt was obtained through manual expression of reproductive males. Milt was captured by a plastic transfer pipet, mixed with eggs, and allowed to sit for 10 minutes. 1 micron filtered sea water was then added to developing embryos.

### Sea robin larval rearing

Sea robin larvae were raised at 18 °C in deep well dish petri dishes for three days in 0.22 micron filtered sea water with 130 μl methylene blue/L. Half water changes were done daily. Prior to the hatching, embryos were transported from MBL to Roger Williams University (RWU) in 50 mL conical tubes. At RWU, they were placed in recirculating Modular Larval Rearing Systems (MoLaRS)^38^. To optimize hatching conditions, a salinity of 36 ppt was maintained, with the ambient water temperature consistently between 20-22 °C. Larvae hatched on day 4 as yolk sac larvae, and initiated first feeding on day five. MoLaRS were pre-stocked with *Parvocalanus sp*. copepods (nauplii) after hatching. As larvae developed, salinity was gradually lowered to lab ambient (30 ppt), and maintained between 30-32 ppt during the course of larval development. *Tisochrysis galbana* supplemented with *Tetraselmis sp*. and/or *Nannochloropis sp.* served to provide "greenwater" background contrast, enhanced feeding response, and provided food for copepods. The copepod stocking density was maintained at 2 to 4 per mL for both nauplii and adults. 10-15 days post-hatch, larvae were supplemented with *Pseudodiaptomus pelagicus*. Larvae began setting after approximately 3 weeks and juveniles were fed enriched *Artemia salina* meta-nauplii. Instar 2 *Artemia* were enriched with LARVIVA Multigain (BioMar SAS) at concentration of 0.5 grams per 1 L of seawater for 4 hours and cold stored (4 °C) until feeding. Five weeks post-fertilization, juveniles were transitioned onto B2 TDO Chroma Boost pellets (Reef Nutrition) and fed larger pellets as they grew. E-series sand (Holliston Sand Co Inc, Burrillville, Rhode Island, USA) was introduced to the tanks to prevent potential fin abrasion.

### Genome generation

Liver tissue from a female *Prionotus carolinus* and a female *Prionotus evolans* was dissected, flash frozen, and sent to Dovetail genomics for genome generation and assembly. DNA was extracted using a Qiagen Genomic DNA extraction kit. DNA samples were quantified using a Qubit 2.0 Fluorometer (Life Technologies, Carlsbad, CA, USA). The PacBio SMRTbell library (∼20kb) for PacBio Sequel was constructed using SMRTbell Express Template Prep Kit 2.0 (PacBio, Menlo Park, CA, USA) using the manufacturer recommended protocol. The library was bound to polymerase using the Sequel II Binding Kit 2.0 (PacBio) and loaded onto PacBio Sequel II. Sequencing was performed on PacBio Sequel II 8M SMRT cells.

PacBio CCS reads were used as an input to Hifiasm1 v0.15.4-r347 with default parameters. Blast results of the Hifiasm output assembly (hifiasm.p_ctg.fa) against the nt database were used as input for blobtools2 v1.1.1 and scaffolds identified as possible contamination were removed from the assembly (filtered.asm.cns.fa). Finally, purge_dups3 v1.2.5 was used to remove haplotigs and contig overlaps (purged.fa).

For each Dovetail Omni-C library, chromatin was fixed in place with formaldehyde in the nucleus. Fixed chromatin was digested with DNase I and then extracted, chromatin ends were repaired and ligated to a biotinylated bridge adapter followed by proximity ligation of adapter containing ends. After proximity ligation, crosslinks were reversed and the DNA purified. Purified DNA was treated to remove biotin that was not internal to ligated fragments. Sequencing libraries were generated using NEBNext Ultra enzymes and Illumina-compatible adapters. Biotin-containing fragments were isolated using streptavidin beads before PCR enrichment of each library. The library was sequenced on an Illumina HiSeqX platform to produce ∼30x sequence coverage.

The input *de novo* assembly and Dovetail OmniC library reads were used as input data for HiRise, a software pipeline designed specifically for using proximity ligation data to scaffold genome assemblies^39^. Dovetail OmniC library sequences were aligned to the draft input assembly using bwa (https://github.com/lh3/bwa). The separations of Dovetail OmniC read pairs mapped within draft scaffolds were analyzed by HiRise to produce a likelihood model for genomic distance between read pairs, and the model was used to identify and break putative misjoins, to score prospective joins, and make joins above a threshold.

### Genome annotation

Liftoff^40^ was used to lift over annotations from the haploid *Gasterosteus aculeatus* genome (NCBI RefSeq Assembly GCF_016920845.1) to haploid *P. carolinus* and *P. evolans* genomes using minimap2 parameters -mm2_options="-r 2k -z 5000".

### CRISPR-Cas9 genome editing

Single guide RNAs (sgRNAs) were designed in Geneious prime software using the Find CRISPR Sites tool. sgRNAs were generated using previously described methods^29^. Table S1 lists all sgRNAs used. The CRISPR-Cas9 mixture consisted of a final concentration of 300 ng/μl sgRNA(s), 1 μg/ul Cas9-2NLS protein (QB3 MacroLab, University of California - Berkeley), 0.05% phenol red, and 70 kDA fluorescein isothiocyanate-dextran diluted 1:5 in the final mixture. 2 M KCl and water were added to a final concentration of 300 mM KCl.

CRISPR-Cas9 reagents were introduced into sea robin embryos using a microinjection approach at the 1-4 cell stage. In brief, quartz needles (Sutter Instrument, QF100-70-10) were pulled using a P2000 micropipette puller (Sutter Instrument) with the following settings: heat = 600, fil = 4, vel = 60, del = 140, pul = 175. Quartz needles were beveled using a BV10 beveler at 20 degrees for 30 seconds. Embryos were lined up in a 1% low melt agarose mold and injected using a Sutter XenoWorks Digital injector.

### Crispant genotyping

Tissues were preserved for genotyping either by fixation in 70% EtOH, or by flash freezing. DNA was extracted using either a DNeasy blood and tissue kit (Qiagen, 69506) or an Allprep DNA/RNA micro kit (Qiagen, 80284). 2x Phusion Green Hot Start II High-Fidelity PCR Master Mix (ThermoFisher Scientific, F566) was used to amplify sequences targeted by the sgRNAs. Products were PCR purified using a QIAquick PCR purification kit (Qiagen, 28106) and quantified using a Qubit DNA High Sensitivity kit (invitrogen, Q32854). Samples were sent for Amplicon-EZ sequencing at Azenta Life Sciences (2×250bp). The raw Illumina reads were checked for adapters and quality via FastQC. The raw Illumina sequence reads were trimmed of their adapters and nucleotides with poor quality using Trimmomatic v. 0.36. Paired sequence reads were then merged to form a single sequence if the forward and reverse reads were able to overlap. The merged reads were aligned to the reference sequence and variant detection was performed using AZENTA proprietary Amplicon-EZ program. Table S1 lists all primers used for generating amplicons. The genotyping results were as follows: 81% percent average mutant reads (*slc24a5* crispants, N = 20) and 1.58% mutant reads (controls, N = 20). 99%, 100%, and 100% average mutant reads (*hoxd12a, hoxa13b, hoxa13a* crispants, respectively. N = 3.) 3%, 1%, 1% mutant reads (controls for *hoxd12a*, *hoxa13a*, *hoxa13a*, N = 1). 86% average mutant reads (*P. carolinus tbx3a* crispants, N = 21) and 0.5% average mutant reads (controls, N = 15). 82% average mutant reads (*P. evolans tbx3a* crispants, N = 7) and 1% average mutant reads (controls, N = 6).

### RNA-sequencing and analysis

Juvenile fish were euthanized by immersion in Tricaine-S (Syndel) until 10 minutes past cessation of opercular movements. Tissues were dissected using microscissors and stored in either RNA later (Invitrogen, AM7020) or flash frozen. Tissues dissected into RNAlater were stored overnight at 4 °C and then stored at -20 °C until extraction. Flash frozen tissues were dissected into 1x PBS, liquid was aspirated, and samples were snap frozen in liquid nitrogen and stored at -80 °C until extraction. Libraries were prepared either in house or by Azenta Life sciences. For in-house preparations, RNA extraction was performed using either an RNeasy micro kit (Qiagen, 74034) or an Allprep RNA/DNA extraction kit (Qiagen, 80284). RNA concentration was measured by a Qubit RNA High Sensitivity Kit (Invitrogen, Q32855) and RNA integrity (RIN) was checked by Bioanalyzer at the Stanford Protein and Nucleic Acid Facility (if feasible given low input concentrations). RNA input was normalized across samples and RNA libraries were prepared utilized the SMART-Seq® v4 Ultra® Low Input RNA Kit for Sequencing (Takara, 634890, 634888) followed by the Nextera XT DNA library preparation kit (Illumina, FC-131-1024, FC-131-1096) per manufacturer instructions. Samples were randomized to prevent batch effects. Sequencing was performed by Novogene on either a HiSeq 4000 or a Novoseq 6000 lane (2×150 bp).

For samples prepared by Azenta, frozen tissues were homogenized using a motorized handheld homogenizor and plastic pestles in low volumes of Trizol and flash frozen in liquid nitrogen. Randomization was performed to prevent batch effects. RNA extraction, sample QC, library preparations, sequencing reactions, and initial bioinformatic analysis were conducted at GENEWIZ/Azenta Life Sciences LLC (South Plainfield, NJ, USA). Total RNA was extracted from cells using the Qiagen RNeasy Plus Universal Micro Kit following manufacturer’s instructions (Qiagen). Extracted RNA samples were quantified using a Qubit 2.0 Fluorometer (Life Technologies) and RNA integrity was checked using Agilent TapeStation 4200 (Agilent Technologies).

The SMARTSeq HT Ultra Low Input Kit was used for full-length cDNA synthesis and amplification (Clontech), and Illumina Nextera XT library was used for sequencing library preparation. Briefly, cDNA was fragmented, and an adaptor was added using Transposase, followed by limited-cycle PCR to enrich and add index to the cDNA fragments. Sequencing libraries were validated using the Agilent TapeStation and quantified by using Qubit Fluorometer as well as by quantitative PCR (KAPA Biosystems). The sequencing libraries were multiplexed and clustered on a flowcell. After clustering, the flowcell was loaded on the Illumina NovaSeq 6000 instrument according to manufacturer’s instructions. The samples were sequenced using a 2×150 Paired End (PE) configuration. Image analysis and base calling were conducted by the HiSeq Control Software (HCS). Raw sequence data (.bcl files) generated from Illumina HiSeq was converted into fastq files and de-multiplexed using Illumina’s bcl2fastq 2.17 Software. One mis-match was allowed for index sequence identification.

For all experiments, RNA-seq analysis was performed in-house. Raw data quality was assessed using FastQC, version 0.11.9^41^. Trimming was performed using Cutadapt version 1.18^42^. Reads were aligned to the annotated *P. carolinus* genome using STAR version 2.7.10b^43^. Read counts per gene were determined using the parameter --GeneCounts in STAR. Differential expression analysis was performed using DESeq2^44^ in R version 4.2.2. N = 6 animals before leg separation and N = 6 animals after leg separation were used in the initial RNA-seq experiment (Fig. 1). N = 8 control animals and N = 10 *tbx3a* crispants were used in the *P. carolinus tbx3a* crispant before leg separation experiment (Fig. 3). N = 4 samples (two *tbx3a* leg samples, one uninjected leg sample, and one uninjected bot sample) were removed due to high sequence duplication levels (visualized in FastQC). N = 6 control animals and N = 8 *tbx3a* crispants were used in the *P. carolinus tbx3a* crispant after leg separation experiment. N = 1 pectoral fin sample from the *tbx3a* crispants was removed before sequencing due to a low RIN score.

### Allele-specific expression analysis

Sequencing quality was confirmed using FastQC, version 0.11.9^41^ and reads were trimmed using Cutadapt version 1.18^42^. Trimmed reads were aligned to a composite *P. carolinus* - *P. evolans* genome using STAR 2.7.9a^43^. Differential gene expression (DGE) between individual species and allele-specific expression (ASE) within hybrids was determined as described previously^45^. For each tissue and stage (e.g. “legs after”), genes with fewer than 10 reads assigned to the *P. carolinus* and *P. evolans* orthologs in each sample were excluded, and each sample was library-size normalized. Genes with significantly different log2 fold-change between *P. carolinus* and *P. evolans* were determined to have differential gene expression (DGE) or allele-specific expression (ASE) (Welch’s t-test; Benjamini-Hochberg FDR < 0.05). Genes were considered to be regulated in *cis-* if the log2 fold-change for both DGE and ASE were significant and in the same direction, and *trans-* if significant only for DGE. Principal component analysis was performed after initial processing with DESeq2^44^ using default parameters. In hybrid samples, size factors were estimated before segregating *P. carolinus* and *P. evolans* alleles.

### Skeletal preparations

Skeletal preparations were performed according to standard protocols^46^. In brief, fish were euthanized and fixed in 10% Neutral buffered formalin (NBF). Samples were rinsed 4x in distilled water overnight while rocking. The next day, fish were stained in Alcian blue, rehydrated in a series of ethanol washes, and rinsed in 30% saturated sodium borate. Samples were incubated in 0.12% porcine trypsin in 30% saturated sodium borate for 3-6 hours at room temperature. Trypsin solution was removed and samples were washed twice in 2% potassium hydroxide (KOH) before being moved into a 0.002% Alizarin red solution in 2% KOH overnight. The next day, samples were bleached with 30% hydrogen peroxide in a 3 part 0.5% KOH and 1 part glycerol solution for up to 8 hours until removal of pigment was sufficient. Samples were then transferred to 100% glycerol with thymol crystals through a 0.5% KOH: glycerol series (1:1, 1:3, 100% glycerol).

### Phenotyping of crispants

Phenotyping of *slc24a5* crispants was performed in the Genome Editing Core at the MBL. Reduction of pigment was scored qualitatively (WT, phenotype) on a Zeiss Discovery V20. Images were taken using Axiovision software. *Tbx3a* and *hoxd12a/hoxa13a/hoxa13b* crispants were phenotyped at RWU and from images of animals taken at RWU. Caudal tails were dissected and placed in 70% EtOH for future genotyping.

### Lobe measurements and analysis

*tbx3a* crispants (N = 4) and control uninjected siblings (N = 5) were euthanized and dissected to expose the dorsal accessory lobes. Standard length of all samples was measured. In FIJI^47^, the freeform tool was used to measure lobe area. Lobe images were scored blinded to genotype. A non-linear regression model in R version 4.2.2 was used to calculate the residuals of the area measurements taking into account the standard length of the fish. Presence of a leg phenotype in the *tbx3a* crispant individuals was observed from images, and lobe measurements of crispants were pooled from the same sides of the body that showed leg phenotypes (N = 2 crispants had unilateral leg phenotypes, N = 1 had bilateral leg phenotypes, and N = 1 crispant had no leg phenotypes and was excluded). Lobe measurement residuals for samples were averaged and a Wilcoxon rank sum test was used to determine significance of the average differences between controls and *tbx3a* crispants.

### Pectoral fin and leg measurements

Adult *P. carolinus* and *P. evolans* were anesthetized using clove oil. For pectoral fin measurements, a flexible tape measure was used to measure the standard length (SL) of the fish in addition to the length of the left and right pectoral fins. The middle of the pectoral fins from N = 10 *P. carolinus* and N = 11 *P. evolans* were measured from the base of the fin to the tip. The pectoral fin length was adjusted according to the standard length of the fish using a linear regression model in R version 4.2.2 and significance of residuals was determined using a Wilcoxon rank sum test, exact = FALSE. For leg measurements, animals were again sedated in clove oil and a flexible tape measure was used to measure the standard length. The leg width of all three legs on the left and right of N = 13 *P. carolinus* and N = 13 *P. evolans* was measured. Leg width was measured at the leg bend, above the shovel structure. Width measurements were regressed against the standard length of the fish in R. A Wilcoxon rank sum test was used to determine significance, exact = FALSE.

### DAPI staining

Whole fish were anesthetized in an overdose of MS-222 and fixed in 4% paraformaldehyde. Tissues (individual legs and/or pectoral fin) were dissected in 1X phosphate buffered saline (PBS) and incubated in solution of 0.5 mg/mL DAPI in PBS for 30-60 minutes in the dark at room temperature with gentle agitation. Tissues were then washed four times in 1X PBS and imaged in 35 mm polystyrene dishes on a AxioZoom v16 microscope (Zeiss). Image analysis was done in FIJI 2. Data were visualized using R version 4.2.2 and Graph pad prism.

### Behavioral analysis

Four groups of six animals (*tbx3a* crispants and wildtype controls, of *P. carolinus* and *P. evolans*) were raised in seawater tanks to ∼2 months. Digging behavior experiments were performed using a behavioral model first tested with adult sea robins. Prior to behavioral experiments, tank sand was cleaned via gravel siphoning and animals were not fed directly before.

Blue mussels (*Mytilus edulis*) were purchased from the local market and dissected into 1-1.5 cm pieces for shallow sand burial. Two PVC-pipe pieces were used to shield visual cues while two mussel pieces were separately buried in the sand. The PVC-pipes were removed, allowing animals to explore the sand. Trials were filmed with GoPro HERO 7 cameras (GoPro, Inc.), outside the behavioral tank. Trials were run for 30 minutes, then the tanks were reset by removing excess mussels and agitating the sand. Ten trials in succession were performed for each tank, ensuring that the mussel pieces changed position in the tank each time.

GoPro videos were uploaded and played back in Windows Media Player to assess capture success. Capture success was defined as a fish locating the mussel, evidenced by mussel consumption or a ‘head snap’ in which fish open their mouths and snap in the direction of the prey item. Each group tank had two mussel pieces or ‘chances of success’ per trial. Data were compiled and visualized in Graphpad Prism.

### Data Availability

All sequencing data will be made publicly available upon publication.

### Code Availability

Code will be made available on Figshare upon publication.

---

**Extended Data Fig. 1:**
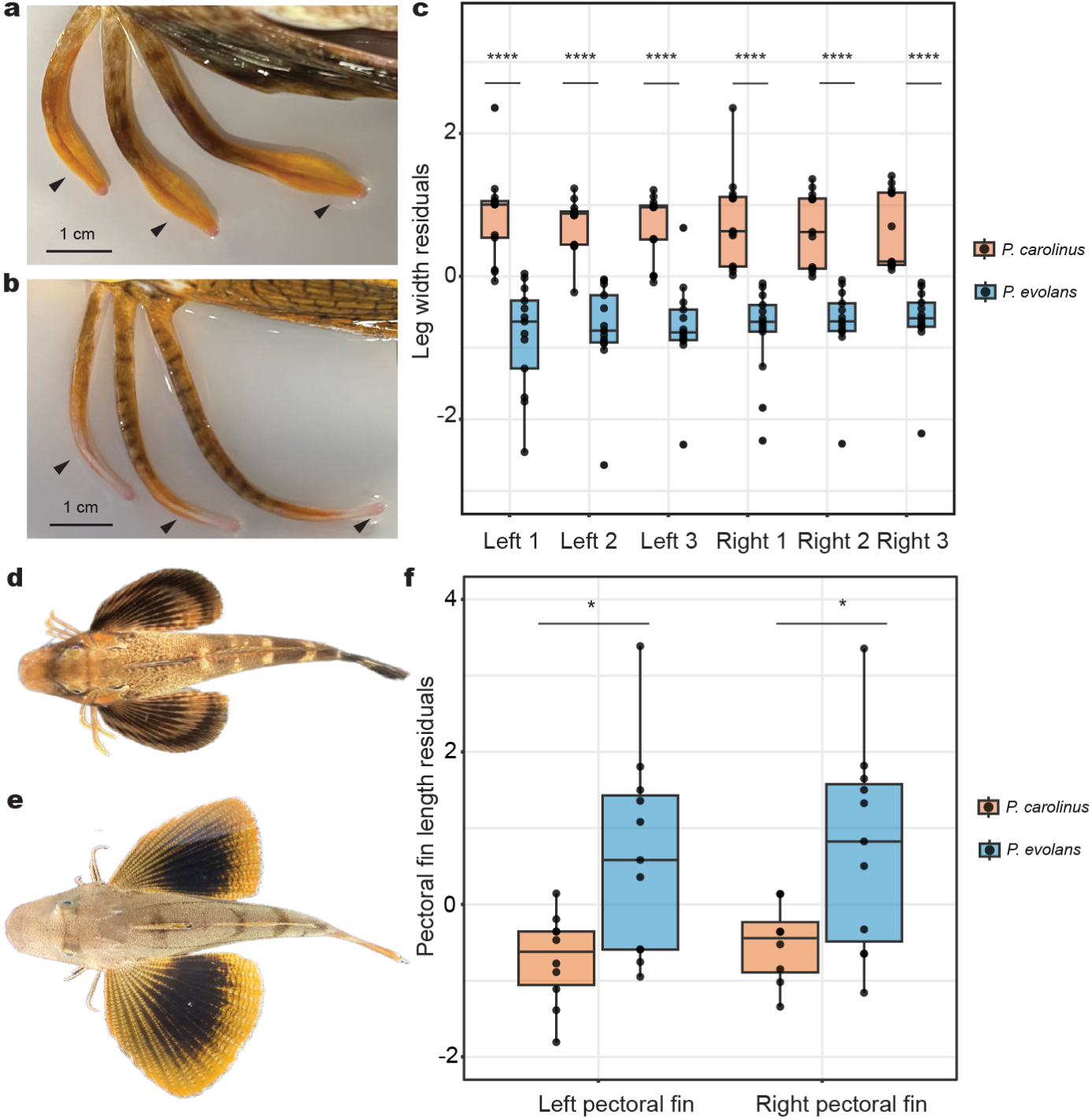
Species-specific leg and fin differences. **a,** *P. carolinus* exhibits thicker legs compared to the legs of *P. evolans* **(b)**, including a shovel-like structure at the tip. **c**, Leg width regressed against the standard length (SL) of the fish. *P. carolinus* legs are significantly wider than *P. evolans* for every leg. **d,** Dorsal view of *P. carolinus* shows pectoral fin span compared to *P. evolans* **(e)**. **f**, Left and right pectoral fin length residuals of *P. evolans* are significantly increased compared to *P. carolinus*. Box and whisker plots show the median at the center line and whiskers in the interquartile range **(c, f)**. Significance determined by Wilcoxon rank sum test. In all graphs, \**P* < 0.05, \*\*\*\**P* < 0.0001. N = 13 *P. carolinus* and N = 13 *P. evolans* (**c**, leg width measurements). N = 10 *P. carolinus* and N = 11 *P. evolans* (**d**, pectoral fin measurements).

**Extended Data Fig. 2:**
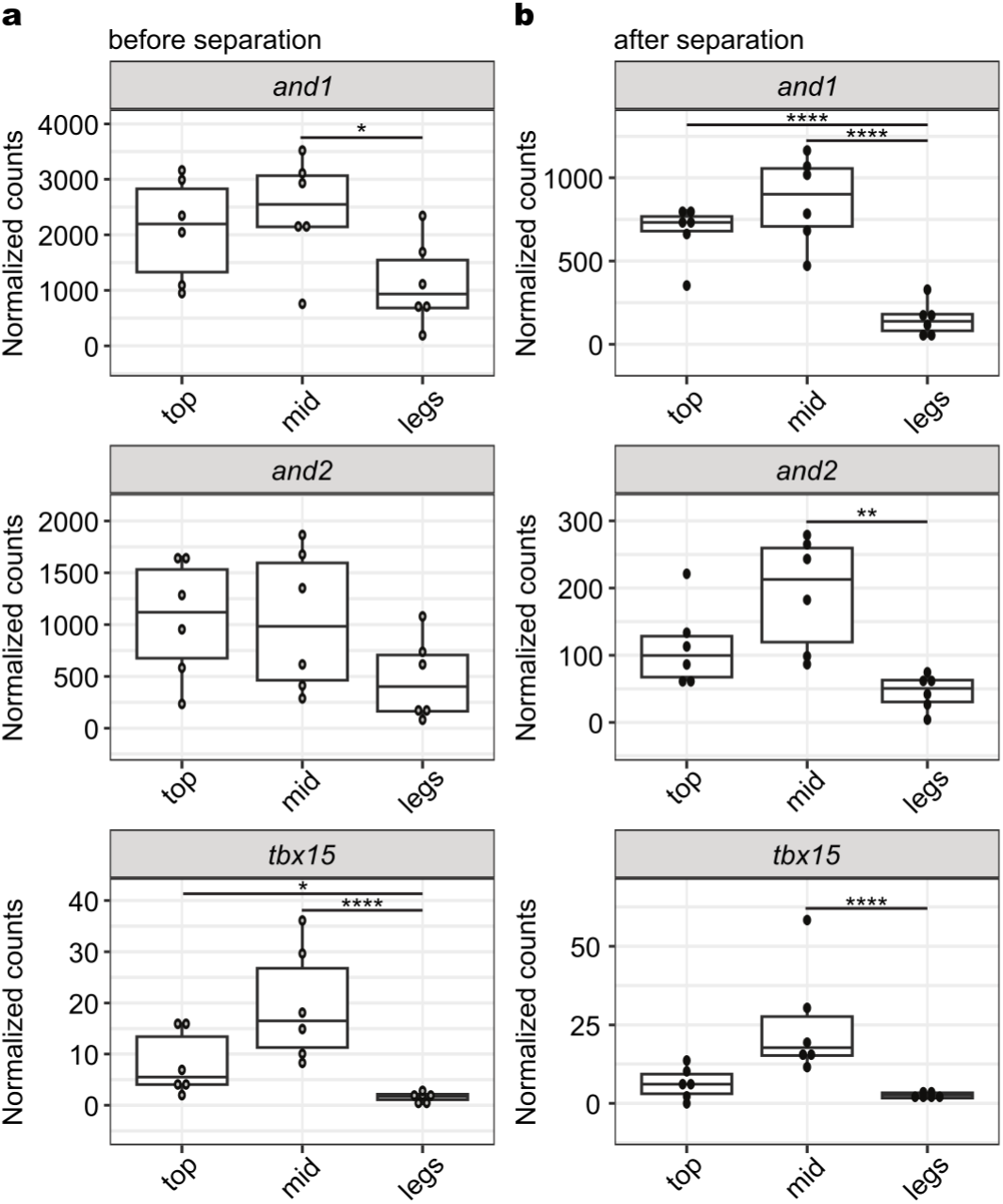
Gene expression in developing fins. **a,** Normalized gene counts from RNA-seq analysis of *and1*, *and2*, and *tbx15* in developing legs and fins, including the top three fin rays (top), middle fin rays (mid) and the legs (leg) before leg separation. **b**, Normalized counts of genes after leg separation. Exact *P-adjusted* values: *padj* = 0.02 (**a**, *and1*), *padj* = 0.02 (**a**, *tbx15* top vs. legs), *padj* = 3.36e-6 (**a,** *tbx15* mid vs. legs), *padj* = 7.68e-12 (**b**, *and1* top vs. legs), *padj* = 9.73e-15 (**b**, *and1* mid vs. legs), *padj* = 0.0018 (**b**, *and2*), *padj* = 1.92e-16 (**b**, *tbx15*). N = 6 animals before separation and N = 6 animals after separation.

**Extended Data Fig. 3:**
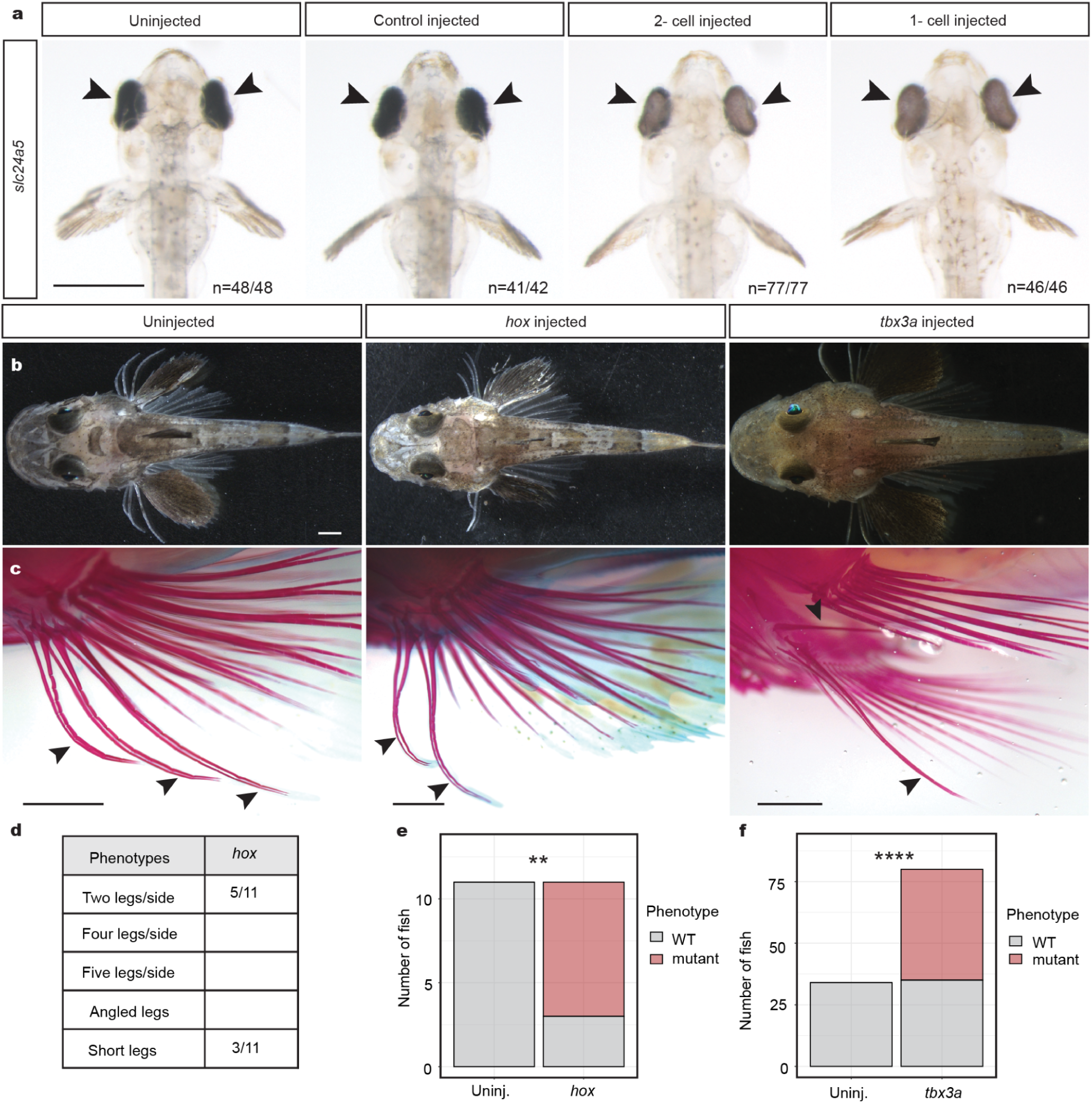
CRISPR-Cas9 genome editing of developmental genes. **a,** Targeting of the pigment gene *slc24a5* at the 1-2 cell stage reduced pigmentation in larval eyes (arrowheads) of injected animals compared to uninjected or control injected siblings. **b**, A dorsal view of uninjected, *hox* injected, and *tbx3a* injected sea robins shows normal gross morphology. Arrowheads point to legs in uninjected juveniles. **c**, Alizarin red staining shows fewer legs in the *hox* injected fish, as well as alteration to leg angle in the *tbx3a* fish. **d**, Quantification of crispant phenotypes in *hox* injected animals. **e**, **f,** Calculation of phenotype significance in *hox* crispants (**e**) and *tbx3a* crispants (**f**). Fisher’s exact test used for calculating significance. Exact *P-values: P* = 0.001 (**e**), *P* = 5.29e-10 (**f**). All scale bars = 1 mm.

**Extended Data Fig. 4:**
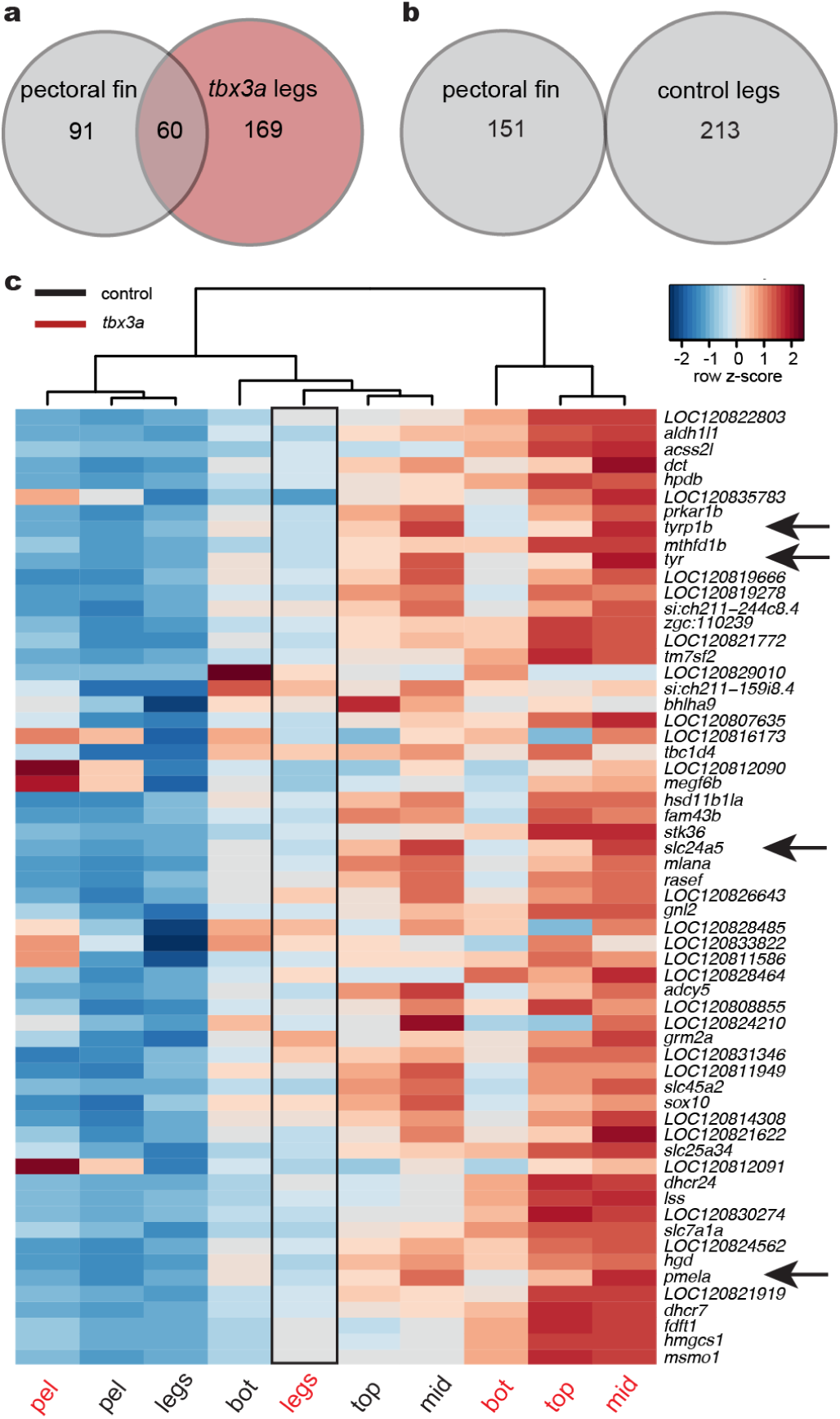
Gene expression in *tbx3a* crispants before separation. **a,** Venn diagram of overlapping genes upregulated in pectoral fins compared to control legs and *tbx3a* crispant legs compared to control legs (*padj* < 0.1). **b**, Venn diagram showing no overlap between genes upregulated in pectoral fins and genes upregulated in control legs compared to *tbx3a* crispant legs (*padj* < 0.1). **c**, Heatmap of the 60 intersecting genes identified in (**a)**. Arrows point to pigment genes upregulated in *tbx3a* crispant legs compared to control legs. Exact *P-adjusted* values: *padj* = 0.03 (**c**, *tyr, tbx3a* crispant legs vs. control legs), *padj* = 0.03 (**c**, *tyrp1b, tbx3a* crispant legs vs. control legs), *padj* = 0.01 (**c**, *slc24a5 tbx3a* crispant legs vs. control legs), *padj* = 0.07 (**c**, *pmela, tbx3a* crispant legs vs. control legs).

**Extended Data Fig. 5:**
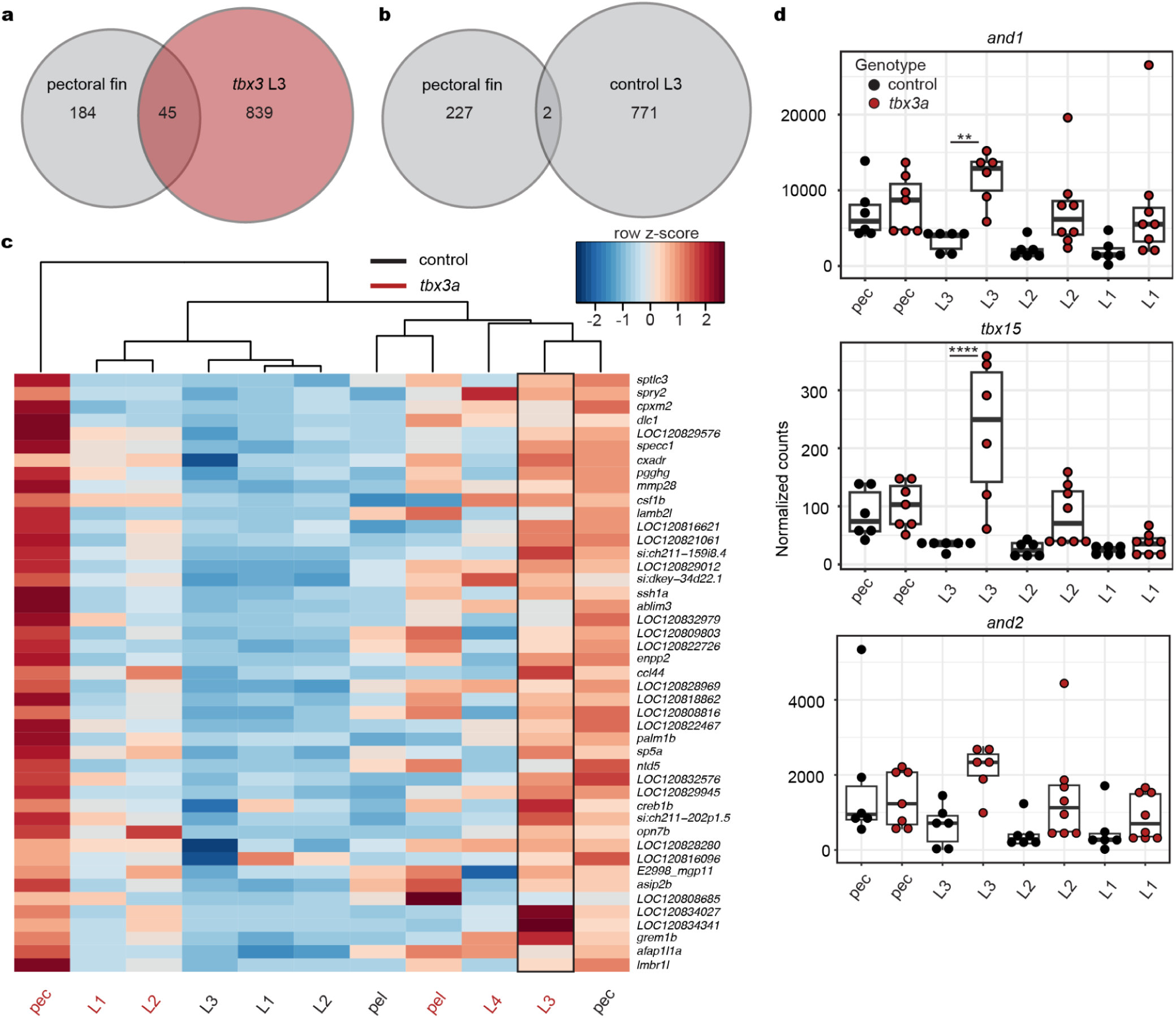
Gene expression in *tbx3a* crispants after separation. **a,** A diagram shows the tissues used in RNA-seq, including individual legs, the entire pectoral fin, and entire pelvic fin. **b,** Venn diagram of 45 overlapping genes upregulated in the pectoral fin compared to control leg 3 and *tbx3a* crispant leg 3 compared to control legs (cutoff set at *padj* < 0.1). **c**, Venn diagram showing two genes are upregulated in pectoral fins and in control legs compared to *tbx3a* crispant leg 3 (cut off set at *padj* < 0.1). **d**, Heatmap of the 45 intersecting genes identified in (**b)**. **e**, Normalized gene counts of *and1* and *tbx15. And1* and *tbx15* expression is upregulated in *tbx3a* crispant leg 3 compared to control leg 3. *And2* counts show similar trends but do not rise to significance. Exact *P-adjusted* values: *padj* = 0.007 (**e**, *and1*, *tbx3a* crispant leg 3 vs. control leg 3), *padj* = 2.27e-9 (**e**, *tbx15*, *tbx3a* crispant leg 3 vs. control leg 3). N = 6 control animals and N = 8 *tbx3a* crispants (N = 7 *tbx3a* pec fins). N = 2/8 crispant animals had two legs.

**Extended Data Fig. 6:**
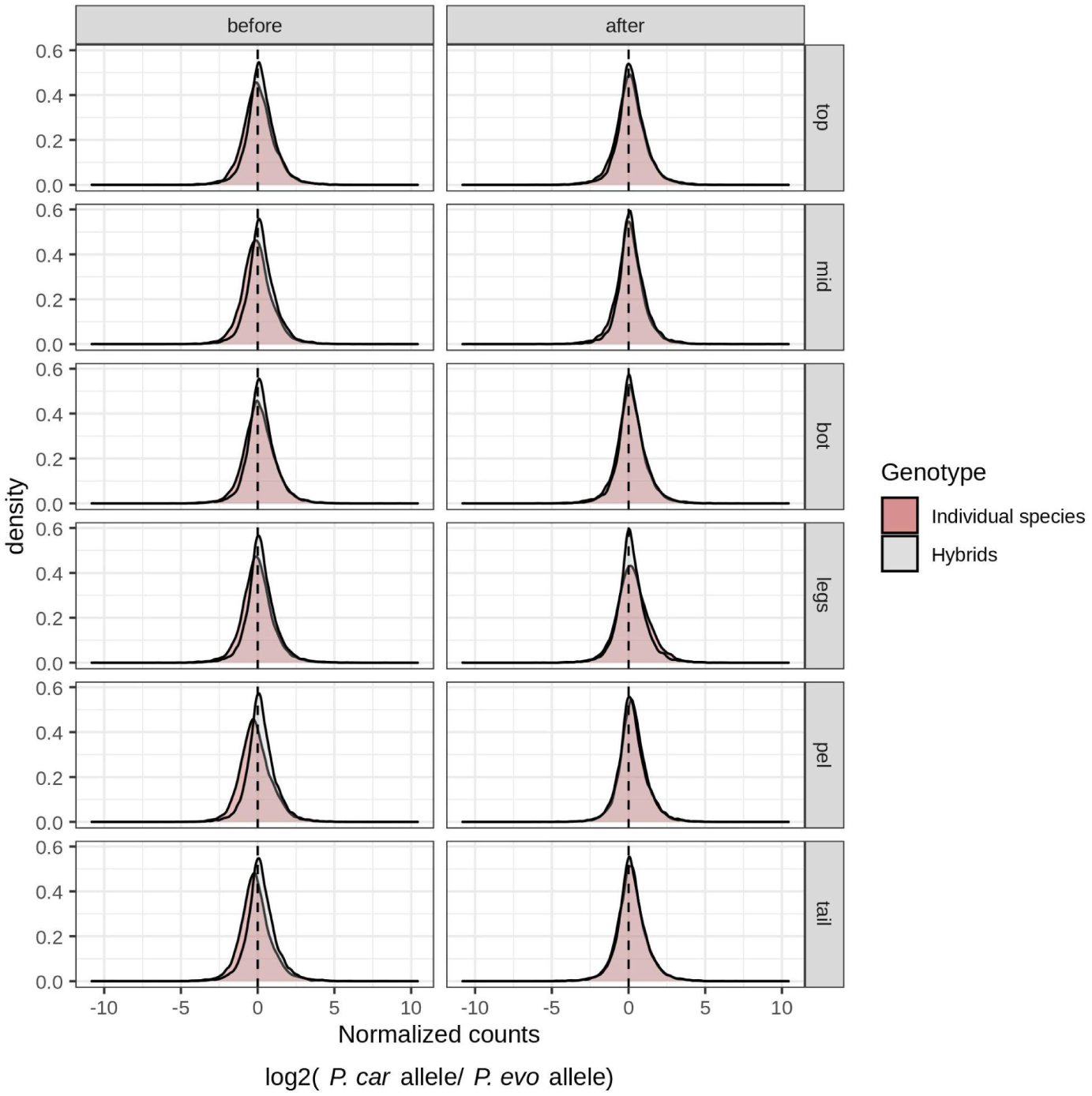
Species-specific differential gene expression. Density plots of normalized counts for *P. carolinus* versus *P. evolans* alleles in individual species (gray) and F1 hybrids (red) across different tissues (top, mid, bot, legs, pel, and tail) and developmental stages (before and after leg separation). The distribution of allele-specific expression in F1 hybrids is slightly biased toward *P. carolinus* alleles, which may reflect paternal allele bias or species-specific expression bias as has been noted in other studies^48, 49^.

**Extended Data Fig. 7:**
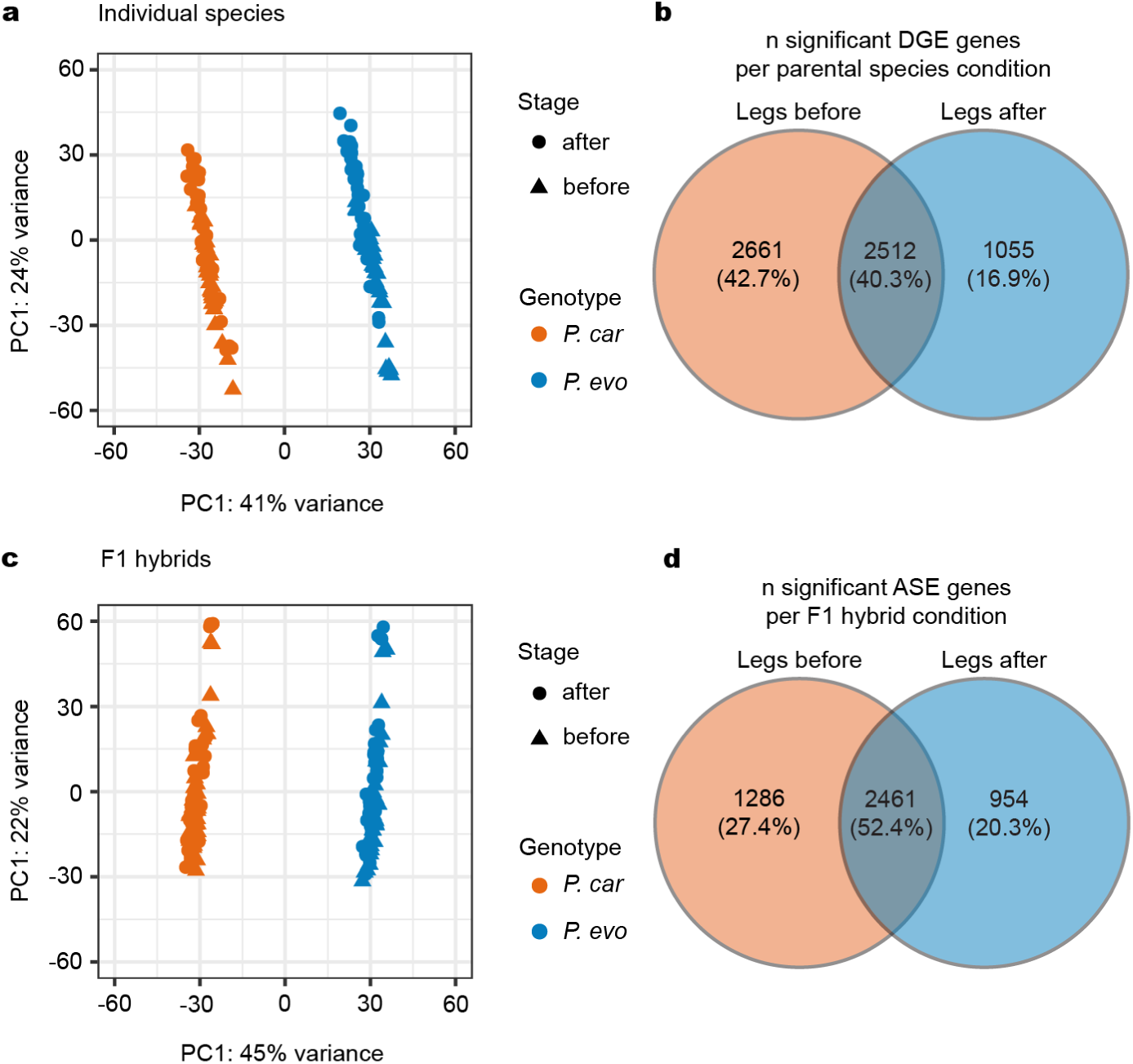
Gene expression in individual species and hybrids. **a**, Principal component analysis of legs before and after separation in *P. carolinus* (*P. car*) and *P. evolans* (*P. evo*). **b**, Venn diagram of the number of genes with significant differential expression between *P. car* and *P. evo* legs before separation (orange) and after separation (blue) in parental species. **c**, Principal component analysis of legs before and after separation in F1 hybrids. **d**, Venn diagram of the number of genes with significant differential expression between *P. car* and *P. evo* alleles in legs before separation (orange) and after separation (blue) in F1 hybrids. Tissues include legs, bot, mid, top, pel, and tail.

**Extended Data Fig. 8:**
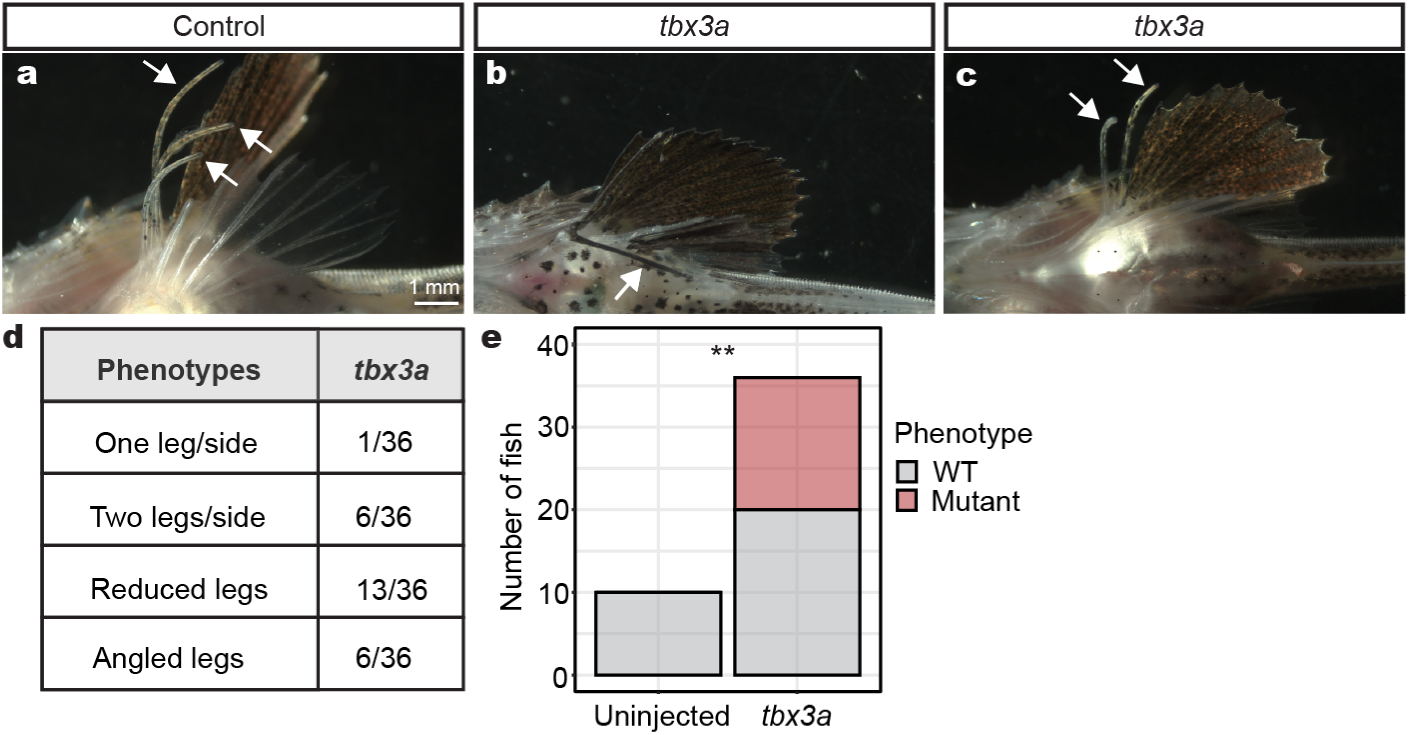
*P. evolans tbx3a* crispant phenotypes. **a,** Control legs in a *P. evolans* juvenile. **b**, An angled leg in a *tbx3a* crispant. **c**, A *tbx3a* crispant with two legs. **d**, Quantification of variable *tbx3a P. evolans* phenotypes. **e**, Calculation of phenotype significance in *tbx3a* crispants. Fisher’s exact test used for calculating significance. Exact *P-value: P* = 0.009. N = 10 controls and N = 36 *tbx3a* crispants.

**Extended Data Fig. 9:**
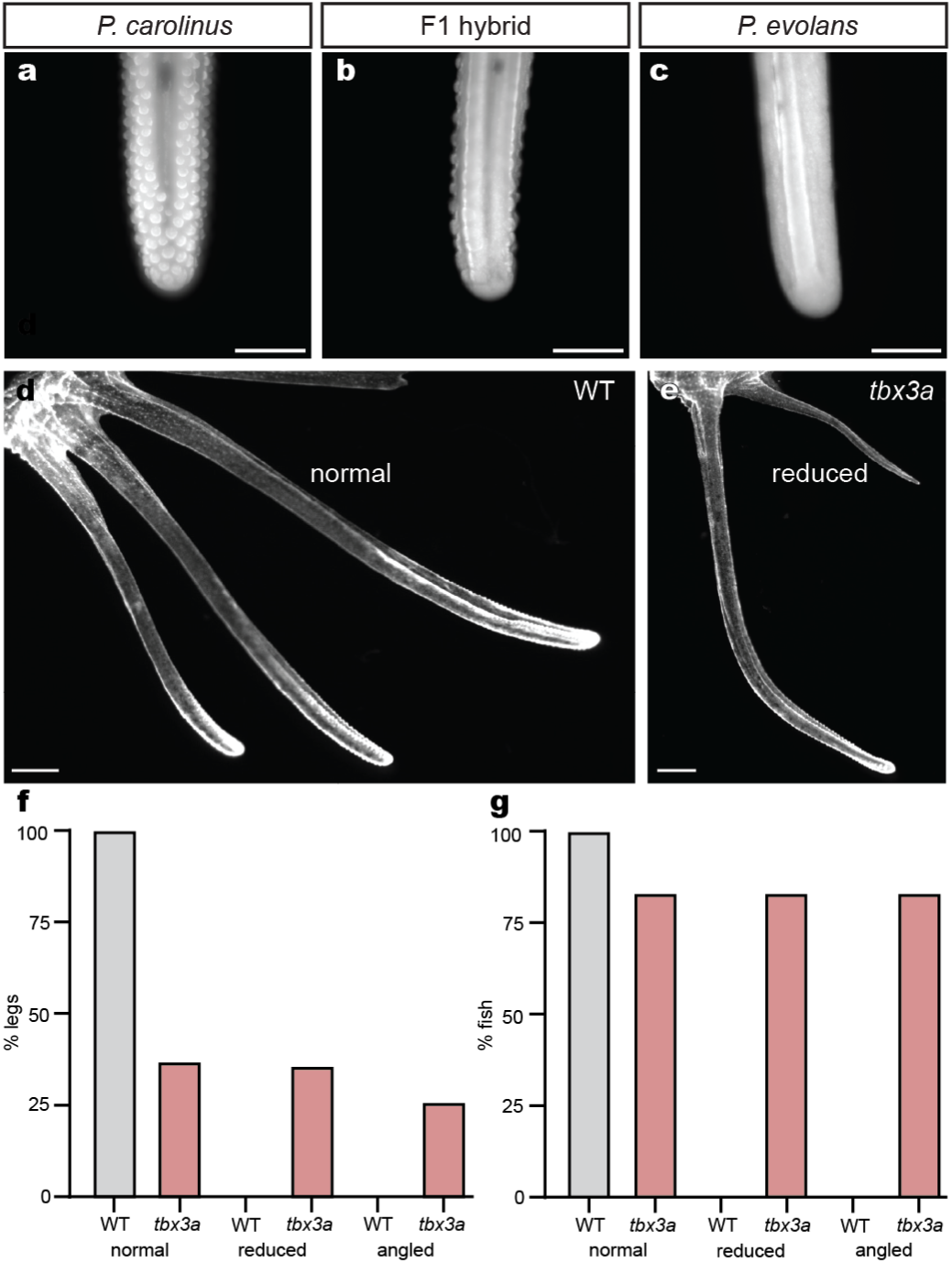
Papillae and leg phenotypes across species, hybrids, and *tbx3a* crispants. **a,** A juvenile *P. carolinus* leg exhibits robust papillae. **b**, An F1 hybrid shows intermediate papillae while a *P. evolans* animal lacks papillae (**c**). **d,** Control *P. carolinus* animals have three normal legs while legs in *tbx3a* crispants can appear normal or reduced in size (**e**). **f**, Quantification of different leg phenotypes in *P. carolinus* control and *tbx3a* crispants. **g**, Quantification of the percentage of fish exhibiting phenotypes. N = 36 control legs and N = 35 *tbx3a* legs analyzed (**c**) and N = 6 control and N = 6 *tbx3a* crispant fish analyzed (**d**). Papillae/mm were measured in the same animals and quantified in Fig. 4e. Scale bars, 500 μm (**a, b, c**); 1 mm (**d**, **e**).

